# Childhood socio-economic disadvantage predicts reduced myelin growth across adolescence and young adulthood

**DOI:** 10.1101/589713

**Authors:** Gabriel Ziegler, Michael Moutoussis, Tobias U. Hauser, Pasco Fearon, Edward T. Bullmore, Ian M. Goodyer, Peter Fonagy, Peter B. Jones, NSPN Consortium, Ulman Lindenberger, Raymond J. Dolan

## Abstract

Socio-economic disadvantage (SED) increases exposure to life stressors. Animal research suggests early life stressors affect later neurodevelopment, including myelin developmental growth. To determine whether human childhood SED affects myelination in adolescence and early adulthood we measured the developmental increase of a sensitive myelin marker, magnetization transfer (MT), in a longitudinal study. Childhood SED was associated with globally reduced MT, as well as slower intra-cortical MT increase in widespread sensory-motor, cingulate, insular and prefrontal areas and subcortical areas. Parental education partially accounted for the SED effects on MT increase, while positive parenting provided a partial protection against the impact of SED. Thus, early socio-economic disadvantage, a vulnerability factor for a range of ill-health outcomes, is a risk factor for aberrant myelin growth during a critical developmental period that is associated with a high risk of psychiatric disorder.

## Introduction

Socio-economic disadvantage (SED) increases exposure to childhood adversity (Pascoe et al., 2016). In turn, adversity is associated with specific patterns of impairments in human brain development (Evans & Cassells, 2014; Johnson, Riis, & Noble, 2016; McDermott et al., 2019; Whittle et al., 2017). The impact of early life socio-economic disadvantage (SED) on brain development is poorly understood at the microstructural level. Here, we make use of a unique longitudinal sample of young people and *in-vivo* quantitative MRI, a measure of macromolecular content sensitive to myelin content, to examine the effects of SED on myelin development.

Animal studies, where early stressors are under experimental control, show a causal impact of adversity on brain growth, even during adolescent development (Howell et al., 2013; Liu et al., 2012; Zhang, 2017). Some experimental manipulations, such as variable foraging demand (Coplan et al., 2016, 2006), bear resemblance to the resource uncertainty encountered in human SED, but it is still difficult to generalize from animal stress to human disadvantage. Thus, mindful of the dangers of making causal implications (Wax, 2017), it is important to map longitudinally human microstructural brain changes such as myelination in relation to SED. This can allow a more detailed comparison with animal findings, and close a causal explanatory gap (Donahue et al., 2018). Cortical myelin likely reflects local neuritic insulation and fibre density (Glasser et al., 2014). It enables myeloarchitectonic parcellation (Glasser et al. 2016) and influences neuronal dynamics (Demirtas et al., 2019).

We recently mapped neurotypical myelin development during adolescence and young adulthood, using myelin-sensitive magnetization transfer saturation (MT) (see also Turati et al., 2015) and showed myelin growth is tied to mental health traits (Ziegler et al., 2018). Human neuroimaging studies, for example in childhood as a function of cortisol reactivity (Sheikh et al., 2014) or in young adulthood as a function of developmental stressors (Jensen et al., 2017), have linked adversity to alterations in white matter myelin. However, these findings beg the question as to how myelination unfolds in detail during adolescence, in both cortex and adjacent white matter, as a function of early socio-economic disadvantage, rather than of specific stressors.

Here we seek to clarify whether patterns of longitudinal myelin growth during late development are associated with early SED as well as establish what role parenting plays in mediating or moderating any relationship (Hair, Hanson, Wolfe, & Pollak, 2015; Noble et al., 2015; Whittle et al., 2017). Under the hypothesis that SED impacts brain development, we predicted that neighbourhood-level indices of deprivation would be associated with both the mean level of myelination, and rate of myelin growth, during late brain development. We found evidence in support of the latter but not the former. Family factors, specifically lower parental education and poorer reported parenting, explained important aspects of the observed relationship.

## Results

### Early disadvantage is associated with slower myelin growth

We obtained 497 repeated structural MRI scans on 288 (51.7% female) healthy participants between 14 and 25 years of age (see Methods). We used an observational accelerated-longitudinal design of these community dwelling English young people and focused on longitudinal findings, a focus which obviated biases seen in cross-sectional estimates with respect to development (Lindenberger, Von Oertzen, Ghisletta, & Hertzog, 2011). We used MT rather than morphometric indices as a measure of (myelin) development, following the priorities highlighted recently by neurodevelopmental researchers (Walhovd, Fjell, Giedd, Dale, & Brown, 2017).

We analysed grey and white matter MT growth related to age, within as well as across participants, using the efficient ‘sandwich estimator’ (Guillaume et al., 2014) and adjusting for curvilinear trajectories (for details cf. methods and Ziegler et al., 2018 focussing on effects of demographics on MT in same sample). This analysis allowed us to index selected longitudinal results in terms of MT growth rate (over follow-up visits) adjusting for the participants’ baseline age.

Our primary measure of SED was the neighbourhood poverty index (NPI), provided by the UK Office of National Statistics (Fry, 2010). NPI was available for all 288 participants at the time of scanning, and for 185 (45.7% female) of these participants before 12 years of age (Supplementary Figure S1). We examined putative explanatory variables both by entering them as covariates and moderating factors in the imaging analyses (cf. methods).

Strikingly, we found that worse early life SED (i.e. living in a more deprived neighbourhood in before age 12) correlated robustly with reduced rate of MT increase (MTr) in multiple brain areas (Figure 1 and Supplementary table 1). This supported an hypothesis that SED is associated with reduced myelin growth (accounting for age and visits) during late adolescent development (Howell et al., 2013; Jensen et al., 2017). Moreover, a reduction in MTr was also observed globally (MTr within whole-brain grey matter, t(321)=−2.87, p=.0022, one-sided, cf. methods). This contrasted with current abode in a disadvantaged neighbourhood, which predicted reduced growth only weakly (Supplementary Figure S2). Furthermore, there were no brain areas where SED was significantly associated with increased MTr (or MTm), which might be expected if some changes reflected adaptive or compensatory early growth (Ono et al., 2008; Ziegler et al., 2018).

**Figure 1.**
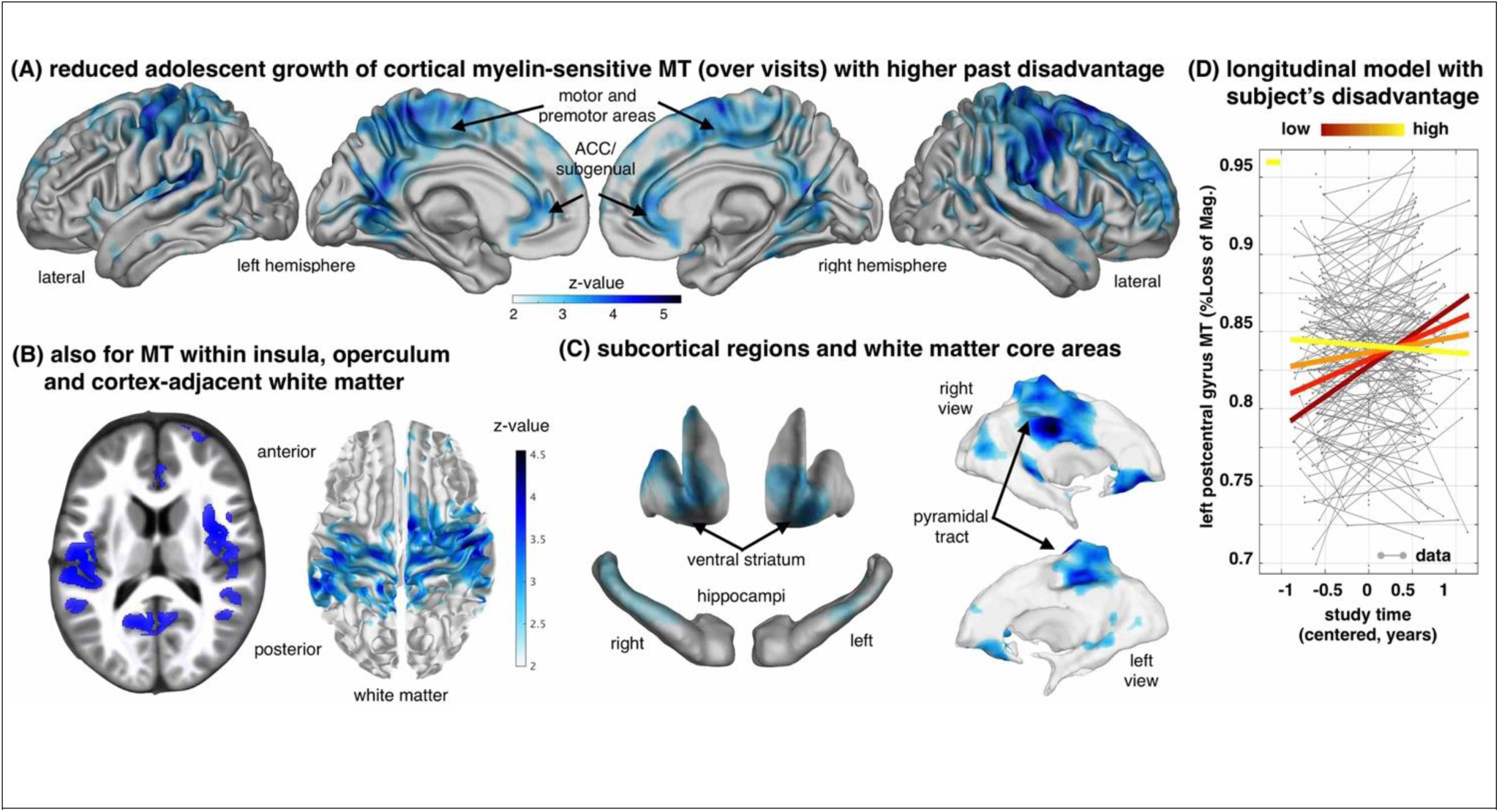
Socio-economic disadvantage (SED) during childhood is associated with slower myelin-related MT growth during coming of age. **A.** Age-typical growth slows down with worsening early life SED in cortex, especially in bilateral precuneus/posterior cingulate, sensory-motor, premotor, sub-genual, and prefrontal areas (z-maps showing negative SED by time/visit interactions, p<.05 FDR corrected, N=328/185 scans/subjects in A-C, 45.7% female). **B.** MT growth is also reduced in insula, operculum (left) and the white matter adjacent to the affected cortex (right). **C.** Hippocampal and striatal grey matter, and core white matter regions also showed reduced growth of MT. **D.** SED-dependent rate of MT growth over visits in a region-of-interest sphere encompassing the central operculum/posterior insula (radius 6mm, centre at x= −47, y=−22, z=13 mm, MNI). Plot shows subjects with higher SED (light yellow) compared to low SED subjects (dark red) express significantly less MT growth over visits (coloured lines in right panel indicate the interaction effect; y-axis: MT; x-axis: time of scan in years relative to each subject’s mean age over visits).

Early life SED correlated with reduced intra-cortical MTr across multiple brain regions including mid- and posterior cingulate, precuneus, operculum, insula (Figure 1a-b) and prefrontal areas, especially on the right. Interestingly, MTr of juxta-cortical white matter and also subcortical regions was similarly reduced (Figure 1b-c, Supplementary table 2). These rate reductions were most pronounced in highly myelinated areas (e.g. M1, S1; for maps see e.g. Glasser et al., 2014) and in regions showing significant developmental MT increase in the very same group of participants (Ziegler et al., 2018).

Our hypothesis that the mean level of myelin, as reflected by the mean MT over visits would show a similar reduction with SED was supported at the global brain level (MTm; accounting for age, visit, sex, and confounds - see methods). Higher SED was associated with reduced global MTm (t=−2.15, p=.016, one-sided, df=321). The local analysis however showed no significant associations between early life SED and MTm (FDR corrected). In the light of the significant global result, greater statistical power may be needed to resolve local MTm effects.

### The contribution of family factors

How early life SED influences brain myelination is likely to involve family-related factors, indexed both by demographic characteristics but also how parents look after their children (Whittle et al., 2017). We examined whether parental education, parental occupation (a proxy for family income), and self-report measures of parenting quality accounted for the effect of SED on myelin growth trajectories (Ronfani et al., 2015; Sarsour et al., 2011). Parental educational qualifications were translated to years-of-education to derive a continuous measure. This variable partially accounted for our MT findings, broadly replicating but also expanding upon existing findings (Noble et al., 2015). Specifically, while peak clusters remained significant, their extent was much reduced, especially in the medial surface of the brain. For example, the left subgenual, right medial motor and right posterior cingulate clusters were largely abolished (cf. Figure 2A vs. Figure 1A). Therefore parental education appears to index influences overlapping with neighbourhood-level SED, but further important influences operate within SED. Against our hypothesis, controlling for poorly paid parental occupation (“Standard Occupational Classification: SOC2000 | HESA,” n.d.) without other family covariates had little impact on the relationship between SED and myelination (Figure 2B).

**Figure 2.**
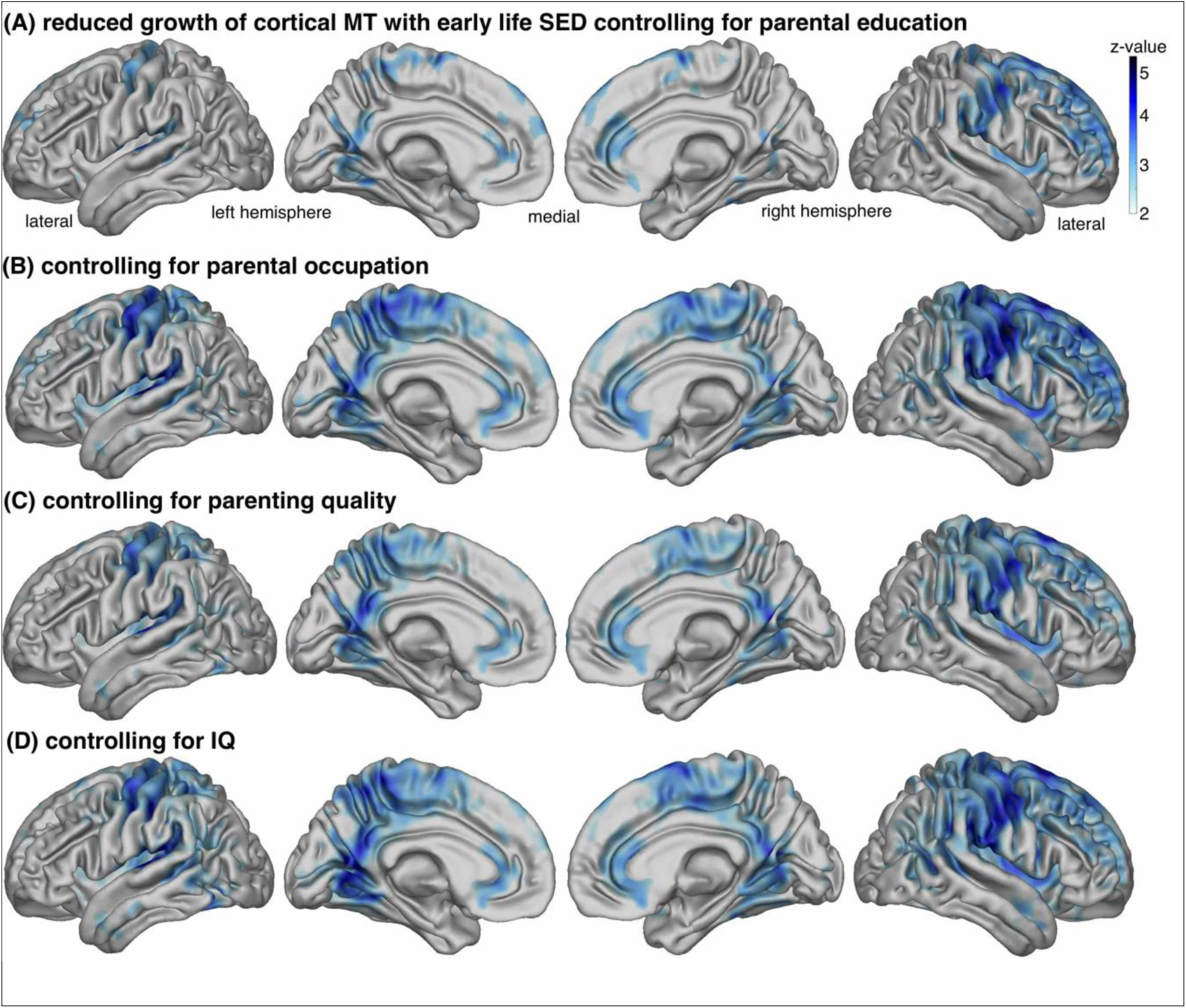
Slower growth of myelin-sensitive MT as early life SED increases is partially explained by parental education but not other factors. We present z-maps showing negative SED by time/visit interactions, p<.05 FDR corrected, N=328/185 scans/subjects, 45.7% female, when additionally controlling for multiple covariates and their respective time/visit interactions in A-C). **A.** Controlling for parental education reduces the impact of SED, in medial motor and premotor areas more than right lateral prefrontal ones (cf. Figure 1A) **B.** In contrast, controlling for parental occupation has minimal impact. **C.** Overall parenting quality has small impact (cf. see Figure 3). **D.** Controlling for time-varying IQ raw scores (here, WASI matrix) has negligible effect on the interaction (similar results for vocabulary, not shown). Controlling for baseline (or mean) IQ over the study period had a very similar effect. Colour scale is identical for A-D.

We next examined family factors proximal to the experience of participants, specifically subjectively reported parenting quality. Here, we used overall parenting quality as an independent variable, estimated by subtracting scores of negative parenting (e.g. harsh parenting or neglect) from those of positive ones (e.g., parental warmth or praise; see Methods). Component negative and positive scores were derived from three self-report questionnaires (Kiddle et al., 2017). We found that parenting quality was not associated with MTr and thus could not mediate the effect of SED on MTr (Figure 2C).

By contrast to the absence of mediation effects, we found a significant moderating effect of parenting on SED, such that better parenting significantly reduced the detrimental effect of early life SED on adolescent MTr. This expands upon existing studies (Sheikh et al., 2014; Whittle et al., 2017). Topographically, this moderating effect was largely confined to lateral prefrontal cortical MT (Figure 3B) and subcortical MT (Supplementary table 3). Thus SED and parental education index overlapping psychobiological influences, while improved parenting quality indexes a separate influence whose presence might have a protective effect in more adverse environments.

**Figure 3.**
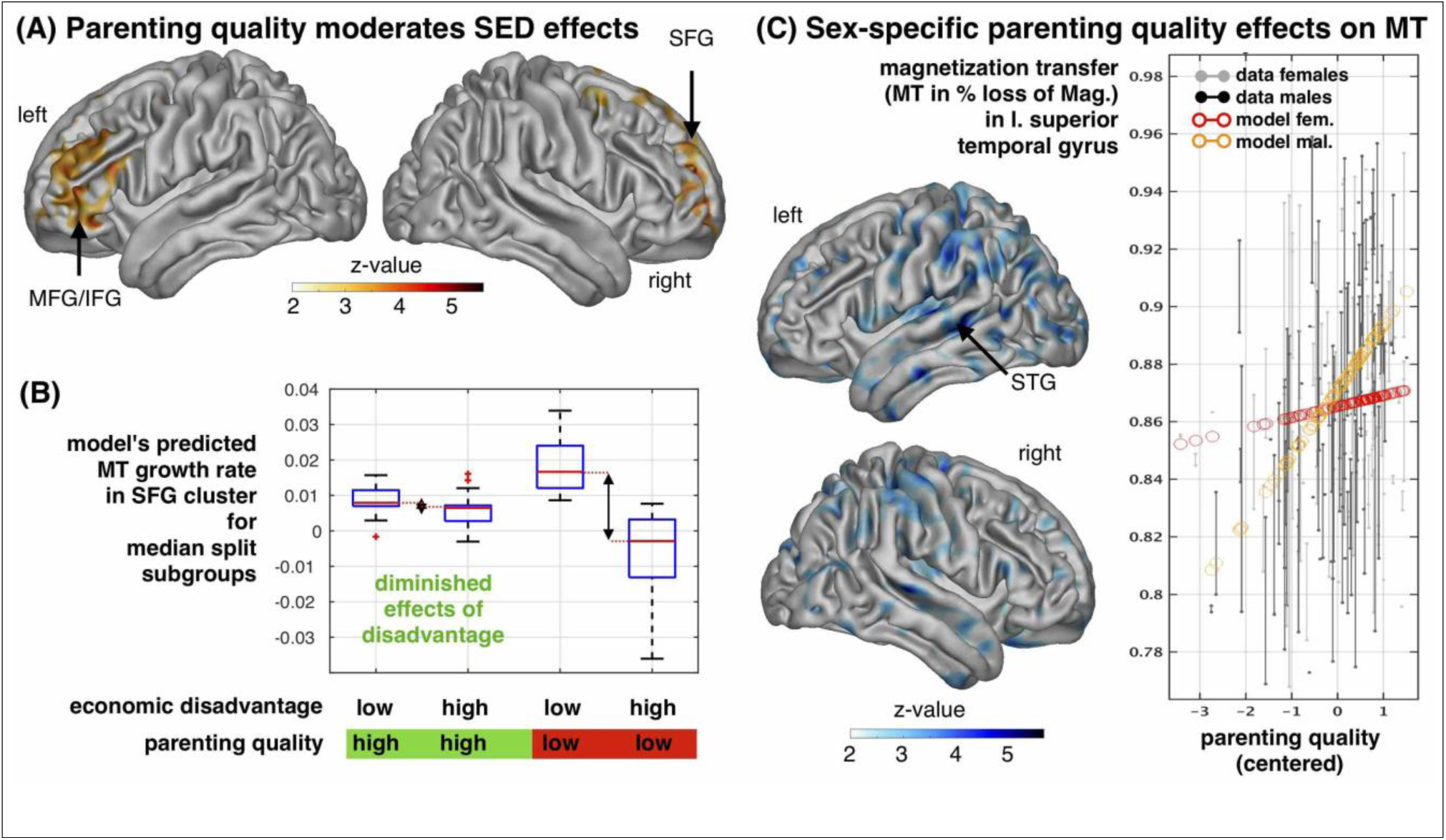
The effect of parenting quality on cortical myelin-sensitive MT and its interaction with SED. **A.** Positive parenting moderated (in the sense of diminishing) the detrimental effect of socio-economic deprivation on prefrontal MT growth. Z-maps show a positive parenting quality by SED by time/visit interaction, p<.05 FDR corrected **B.** Illustration of the moderation effect (A.) within right superior frontal gyrus cluster. We show growth rates (MTr) as predicted by the longitudinal model within median split groups of high vs. low SED and high vs. low parenting quality. The box plot shows averaged data within 6mm sphere around peak voxel illustrating SED by time and parenting by SED by time/visit interaction. Early life SED effects on MTr (illustrated by arrow) are less pronounced in family contexts with high parenting quality. **C.** The effect of parenting quality on MT was significantly steeper in males compared to females (see also Supplementary table 4). Z-maps show negative sex by parenting quality interactions, p<.05 FDR corrected, N=328/185 scans/subjects in A-C, 45.7% female, accounting for age, visit/time, sex, interactions and confounds. The right panel plots MT in superior temporal gyrus (6mm sphere around peak voxel) over parenting quality (x-axis, z-scored) and with adjusted data (grey/black) and model predictions (red/orange, effects of interest: intercept, parenting, sex by parenting). Higher parenting quality only showed a trend towards a positive main effect on cortical MT (p<.001, unc., not shown).

### Individual risk factors

A number of factors measured at the level of the individual might form either important causal contributors, outcomes or markers for the association between SED and MTr. We predicted that lower IQ would be associated with altered patterns of myelination consequent upon SED for three reasons. First, prior evidence suggests that IQ might be related to individual differences of myelination (Dunst, Benedek, Koschutnig, Jauk, & Neubauer, 2014). Second, there is evidence that IQ and socio-economic status share similar genetic determinants (Trzaskowski et al., 2014). Thus, genes that directly contribute to parental socio-economic success may also directly contribute to differences of brain structure, as indexed by IQ. In other words, SED and brain growth could be associated through horizontal genetic pleiotropy (Supplemental Figure S3). Third, the correlation between socio-economic indices and IQ might be explained by morphometric brain measures (McDermott et al., 2019). To test the hypothesis that IQ would index biological processes largely overlapping with those present in SED, we controlled for verbal and matrix IQ scores (Figure 2D), both separately and as a total score. However, accounting for IQ in any of these ways left the relationship between SED and slower MTr unchanged.

We also examined the effects of self-reported ethnicity and alcohol drinking, as these are thought to be reflected in brain structure and connectivity (Noble et al., 2015; Smith et al., 2015). These failed to account for our key findings (Supplementary Figure S4A-B).

We were interested whether determinants of poor physical condition, such as sedentary habits and poor quality nutrition, at least as indexed by body mass index (BMI), explained why more deprived children had lower MTr during adolescence. Thus, we tested a prediction that an association of MT trajectories with early life SED would be partially accounted by an increased BMI. This is important as both SED and poor parenting (Sleddens, Gerards, Thijs, Vries, & Kremers, 2011)) increase the risk of being overweight (Salmasi & Celidoni, 2017), which is in turn associated with deviant white matter development (Kullmann, Schweizer, Veit, Fritsche, & Preissl, 2015).

Mean BMI was positively associated with early SED and negatively to MTm but not MTr, and did it account for the relationship between SED and MTr. As expected BMI increased with age during adolescence, but, importantly, it increased faster the greater the degree of early SED (Figure 4A). Correcting for age, and variables of no interest (see Methods), greater BMI was associated with lower MTm in anterior insula, anterior cingulate and other areas (Figure 4B and 4C). The relationship was, significantly more pronounced in males.

**Figure 4.**
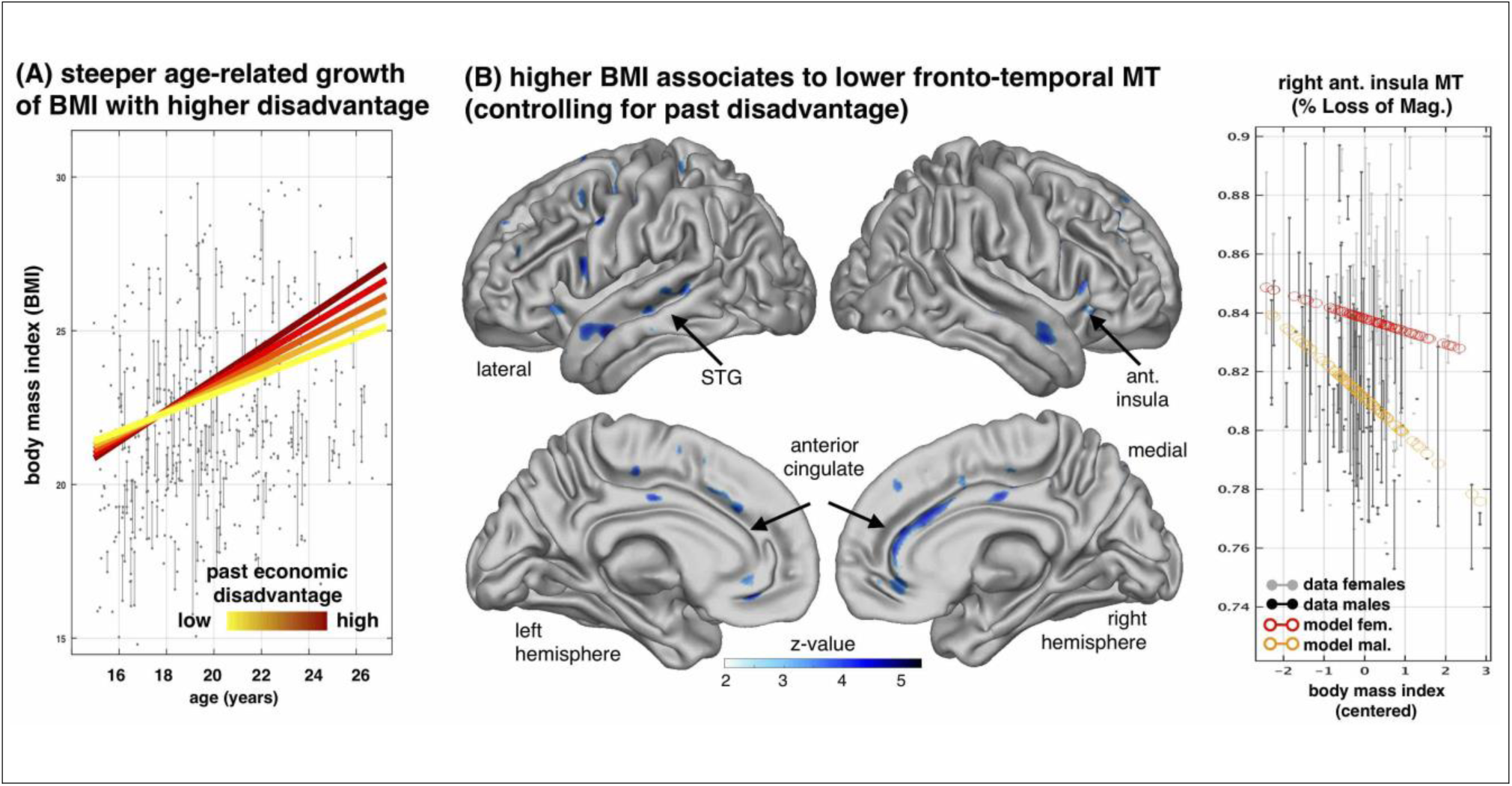
The effect of body mass index on myelin growth. **A.** Early life SED is associated with faster gain of body mass index (BMI) during youth. Linear-mixed modelling revealed positive age effects on BMI (t=4.6, p=3.9e-6, two-tailed, df=559), positive main effects of early life SED on BMI (t=2.05, p=.0407, two-tailed, df=559, N=568/384 observations/subjects) and a steeper age-related increase with higher SED (t=2.2, p=.0265, df=559, two-tailed), accounting for age, visits, sex, and interactions. **B.** Greater BMI is associated with lower cortical myelin-sensitive MT in the anterior cingulate, superior temporal, anterior insula cortex (z-maps showing BMI effects, p<.05 FDR corrected, N=277/155 scans/subjects, 47.5% female, accounting for age, visit/time, sex, early life SED, interactions and confounds). **C.** Right panel shows the plot of MT in insula (6mm sphere around peak voxel) over BMI (x-axis, centred) and with adjusted data (grey/black) and model predictions for sexes (red/orange, effects of interest: intercept, BMI, sex by BMI). The decline of MT with higher BMI is steeper in males than females. See also Supplementary Table 5.

Lastly, we found no significant correlations between SED and global morphometric measures reported in large published studies (Johnson et al., 2016) (Figure S5).

## Discussion

We studied the relationship of early socio-economic disadvantage to myelin growth during adolescence, de facto a relationship between economic disadvantage and deviant neurodevelopment (Johnson et al., 2016). Early disadvantage correlated with a globally reduced myelin and reductions in myelin growth longitudinally, as indexed by the sensitive marker magnetization transfer saturation, MT. Reminiscent of other developmental findings (Sheikh et al., 2014; Whittle et al., 2017), parental education, but not other predictors, accounted statistically for much of this effect. Better parenting moderated the relationship, lessening the effect of economic disadvantage. As hypothesized, increased BMI also indexed deficits in myelin development. However, although BMI was robustly related to SED, BMI had an effect on myelination independently of SED (and not explaining it). These findings have important implications for further, translational research and are relevant to protecting brain development during youth as we shall see.

As our sample was healthy, slower myelination constituted neurodiversity rather than neuropathology. Diversity can be seen as biologically encoding prior beliefs (Dayan, 2012; Moutoussis et al., 2018) conferring adaptation, maladaptation (Jensen et al., 2017) or both, depending on functions yet to be studied. On the one hand, the fact that we observed only reductions in MTr, that these were very extensive, and that they were related to early but not current SED, suggest maladaptation, at least with respect to current circumstances.

Parental education substantially mediated the effects of SED (Figure 2), suggesting these two processes index overlapping biological pathways, reducing MTr. Remarkably, according to our results this is unlikely to be due to IQ-related genes, to parenting quality as perceived by the young person, over-weight, alcohol drinking or ethnicity. It is thus important to further understand what (non-IQ) genes or proximal environmental drivers underpin different myelination trajectories. These may be diverse and could include different parental behaviours not captured by our measure of parenting, the provision of enriched primary schooling, other family environment factors such as parental conflict, or factors associated with early peer processes.

Early poverty is associated with lower life achievement and IQ (Evans & Cassells, 2014) and brain structure may mediate this at least in part (McDermott et al., 2019). In our study, however, baseline IQ scores did not account for the effect of SED. We note that previous research often directly incorporated parental education as a measure of SED, and this may itself strengthen the relation between IQ and SED. Our findings imply that genes directly contributing to both (neighbourhood) SED and MT growth do not underpin our IQ measures. However, SED-dependent vertical pleiotropy may operate. Here, genetic or indeed environmental antecedents would give rise to an intermediate phenotype, on which SED influences would operate to affect neurodevelopment. In this case SED would disproportionally affect the genetically vulnerable (Gage, Smith, & Munafo, 2016). Vertical pleiotropy is further explained in Figure S3.

Primate experiments show that early stressors (Howell et al., 2013) cause long-term problems, including an impact on brain myelination. By analogy, early developmental stressors are likely to be commoner among disadvantaged children. However here, unlike in animal studies, diffuse low-impact mechanisms rather than focal, high-impact ones are more likely to account for the relationship between SED and myelination. This is because the sample was healthy, and severe adversity was under-represented (Figure S1). If broad low-impact influences operate, future research should also prioritize broad influences and interventions over searches for focal, high-risk subgroups.

In primate variable foraging demand (VFD) experiments, it is resource insecurity (Coplan et al., 2006), rather than the average resource level, that affects infant neurodevelopment, suggesting that neighbourhood SED may index stressful economic insecurity for all, not just the poor, consistent with parental occupation not accounting for the effect of SED. That VFD effects may be mediated by maternal preoccupation by insecurity (Coplan et al., 2006) would be consistent with positive parenting modestly mitigating the effects of SED (Figure 3).

We did not replicate a number of findings in the literature connecting SED to macroscopic measures such as grey matter volume or surface area. Our smaller sample, though a limitation, is likely to mean that the myelination effects are more prominent, so easier to detect. Results are also consistent with myelination not being straightforwardly reflected in macroscopic measures (Grydeland, Walhovd, Tamnes, Westlye, & Fjell, 2013).

One limitation of our study that future research should address, is that early SED was assessed retrospectively. Secondly, although chronological age is important, future studies should use endocrinologically precise pubertal stage as a developmental independent variable. Thirdly, studies should address more deprived populations in an international scale. SED influences on myelination may be studied in ‘natural experiments’, such as the precipitous, decade-long exposure to serious SED of Greek children born since 2010 across genetic, educational and occupational strata, resembling a powerful VFD manipulation.

Our research informs future research directions to knowledge about causation and to policies that can protect brain development in young people. First, the functional consequences of the slower myelination must be demonstrated. For example, adverse effects of high BMI on mid-life cognitive function may be restricted to disadvantaged groups (Cohen-Manheim et al., 2017). How SED dependent myelination longitudinally impacts on IQ and on specific sensory-motor, emotional and cognitive functions (Sheehy-Skeffington & Rea, 2017) remains unclear and deserves research attention. Most importantly, our findings indicate that intervention studies aiming to reduce SED and/or improve parental education and parenting in richer countries should prospectively examine their impact on myelination.

In conclusion, neighbourhood deprivation during development was associated with slower myelin growth markers during adolescence and young adulthood. This was independent of baseline IQ, ethnicity or parental occupation, while parental education statistically explained much of the effect and may give clues about causal mechanisms. Causation, functional consequences of myelination and policy implications provide a fertile context for future investigations.

## Materials and Methods

### Recruitment, demographic and psychological measures

Participants were recruited from the Neuroscience in Psychiatry network participant pool (Kiddle et al., 2017). From this pool, also known as the ‘2K sample’, 300 participants were recruited for the present scanning study. We aimed to exclude all but the most minor psychiatric and neurological symptomatology, and therefore screened by self-report participants to not have current or previous relevant medical histories. We finally analysed 497 available brain scans from 288 healthy (149 female) individuals that passed quality control. In particular, data from 100, 167, and 21 subjects with one, two or three visits per person were available, with mean (standard deviation) follow-up interval of 1.3 (0.32) years between first and last visit.

Self-defined ethnic group was asked about shortly after recruitment and in terms of the following broad classes: White (1858 or 80% of declared ethnicity), Black (3.7%), Asian (8.5%) Mixed (6.0%), Other (2.1%), ‘Prefer not to say’. On the day of scanning, participants also completed the Wechsler Abbreviated Scale Intelligence of (WASI) (Kiddle et al., 2017). As ours was a developmental study, we used the raw subscale scores for vocabulary and matrix IQ and explicitly analysed, and their dependence of age. Unless otherwise stated, IQ measurements refer to the time of the first, ‘baseline’ scan.

The measure of overall parenting quality that we used here was a composite of the Positive Parenting Questionnaire (PPQ), Alabama Parenting Questionnaire (APQ) and Measure of Parenting Style (MOPS). All three were obtained within about a month of the first scan (Kiddle et al., 2017). We took the reversed positive parenting total score from the PPQ and the similarly reversed positive parenting scales from the APQ, the negative parenting scales from the APQ (inconsistent discipline, poor supervision, and corporal punishment) and the negative parenting scores for the MOPS (abuse, control and neglect), which were standardized and summed to make a composite negative parenting scale. The internal consistency of the resulting total score was alpha = .96.

As far as parental education is concerned, the young people in our study reported the highest qualification and occupational level of their parents. This data was obtained for the mother, the father, and if applicable the mother’s partner and the father’s partner. These were converted to an ordinal scale, according to a categorization of educational achievement in England - that is, none, primary school, secondary school - GCSE’s, sixth form - A levels, skills-based trainings, undergraduate education, postgraduate education or higher professional training. We then took as starting score the education level of the female parent (usually biological mother) and compared it with the primary male parent (mother’s partner or biological father, in that order of priority). It was unusual for these to differ by more than one on this ordinal scale. Therefore, if only one parent score was available, we used that for ‘parental education’; otherwise we averaged the two prioritized scores.

### Measures of Socio-Economic Disadvantage

We used the following measures of socio-economic disadvantage. As our central measure, we used the neighbourhood proportion of households below the official poverty income around the participant’s residence (“Small area model-based households in poverty estimates, England and Wales - Office for National Statistics,” n.d.) at the time of first scan. We also used an index of parental education (IPE); and the mother’s and father’s SOC2000 occupational class (“Standard Occupational Classification: SOC2000 | HESA,” n.d.).

### MRI data acquisition and longitudinal preprocessing

Brain scans were acquired using the quantitative MPM protocol (Weiskopf et al., 2013) in 3T Siemens Magnetom TIM Trio systems located in Cambridge and London. Isotropic 1mm MT maps were collected to quantify local myelin changes throughout the brain. Analyses were performed using SPM12 (Wellcome Centre for Human Neuroimaging, London, UK, http://www.fil.ion.ucl.ac.uk/spm), the h-MRI toolbox for SPM (Draganski et al., 2011) [https://github.molgen.mpg.de/hMRI-group/Toolbox], Computational Anatomy toolbox (CAT12, http://www.neuro.uni-jena.de/cat/) and the tools described in more detail below and in the methods section of the preceding paper that focussed on effects of demographics in the same sample (Ziegler et al., 2018).

To assess macromolecular growth during development, we used a longitudinal Voxel-Based Quantification (VBQ) pipeline that follows the following steps (for more details and illustration see Ziegler et al., 2018). First, images were serially registered. Each baseline - follow-up mid-point image was then segmented into grey matter (GM), white matter (WM) and cerebrospinal fluid (using the CAT12 toolbox of SPM). MT maps from all time-points were then normalized to MNI space, manually inspected and checked for outliers. A during scan motion proxy was used to discard 10% of images with strongest motion-induced artefacts and as a confounding covariate during all analyses. We constructed masks for both grey and adjacent white matter using SPM neuromorphometrics atlas for tissue-specific analysis of MT parameters. Finally, normalized MT maps were processed with a tissue-weighted smoothing procedure (7 mm FWHM).

### Longitudinal MT image analyses

In order to quantify myelin development we took advantage of the observational accelerated longitudinal design. We focused on how the brains change over study time/visit. We compared and contrasted this to how brain structure varied with scan-midpoint age across participants in the study. To do so, we used the Sandwich Estimator (SwE) method for voxel-based longitudinal image analysis (Guillaume et al., 2014, http://www.nisox.org/Software/SwE/). This so-called marginal model describes expected variability as a function of predictors in a design matrix, while additionally accounting for correlations due to repeated measurements and unexplained variations across individuals as an enriched error term.

In our analyses, we focused on factors time/visits and mean age of the individual (over all visits) on MT across the whole brain. To investigate how exposure to poverty was related to brain trajectories and altered growth, we enriched the models by adding a main effect (SED as measured by the NPI, as a predictor of mid-point MT) as well as the interaction of SED with individual MT change over scan sessions (visits or within-subject study time). The latter metric allowed us to assess how myelin growth is associated with SED (e.g. lower myelin growth upon exposure to high SED), whereas the former indicates how SED relates to overall myelination differences across individuals accounting for other covariates, such as visit, mean age, and sex. We a-priori hypothesized reduced levels of myelin and impaired myelin growth with higher SED. The effects of visit, age, sex, and non-linearities (e.g. in terms of age by age and age by time interactions) of age-related trajectories, and for first order interactions among all demographic variables are presented elsewhere (Ziegler et al., 2018). All analyses were carried out with scanning site, total intracranial volume and motion regressors as confounds. More mathematical details on SwE and longitudinal design specification can be found in supplementary information of Ziegler et al., 2018.

We then tested whether the observed associations of SED might be explained by further covariates, by including the latter on the same footing as SED in analyses. All models were tested for indications of effects of sex, IQ, parental education, parental occupation and self-reported ethnicity. We further conducted moderation analysis in terms of indications whether above family or individual factors show a significant interaction with SED either on mean level MT or MT growth over visits. Thus, two-way interaction with SED (e.g. parenting quality by SED) and three-way interaction terms (e.g. parenting quality by SED by time/visit) were included in addition to all main effects, time, age, sex, and their interactions in SwE models of local MT. We controlled for the False Discovery Rate (FDR, p<.05) during corrections for multiple comparisons in all image analyses.

### Linear mixed effects modelling of global MT and BMI

To assess the effects of SED on global MT and on BMI, we used linear mixed-effects modelling (LME, cf. supplementary information Ziegler et al., 2018). We specified corresponding fixed effects design matrix including time, age, sex, SED, and first order interactions while accounting for confounds. Random-effect intercepts were included and proved optimally suited using likelihood ratio tests. T-values of fixed effects coefficients and corresponding (one-sided) p-values were calculated to test for detrimental main effects of SED and time/visit or age interactions. More mathematical notes on LME and longitudinal design specification can be found in supplementary information of Ziegler et al., 2018.

### Macrostructural measures

Finally, to complement the main focus of this study in assessing SED-related correlates of novel, quantitative, myelin-sensitive MT (using VBQ), we also tested for previously reported relationships of SED with conventional metrics, i.e. Voxel-Based and global Surface-based Morphometry (VBM & SBM). For this purpose we used non-linear registration to obtain normalized (grey and white matter) tissue segment maps using both within-and between-subjects modulation. This was followed by Gaussian smoothing (6mm). Moreover, cortical surface reconstructions of all participants’ midpoint was obtained (using CAT Toolbox), and cortical thickness, surface area, gyrification index, and sulcal depth was assessed in native space, resampled to a surface template and smoothed with 12mm.

## Supporting information

Supplement (all figures and tables)

## Funding

The Wellcome Trust funded the ‘Neuroscience in Psychiatry Project’ (NSPN). All NSPN members (table S1) are supported by Wellcome Strategic Award (ref 095844/7/11/Z). Ray Dolan is supported by a Wellcome Investigator Award (ref 098362/Z/12/Z). The Max Planck – UCL Centre for Computational Psychiatry and Ageing is a joint initiative of the Max Planck Society and UCL. M. Moutoussis receives support from the NIHR UCLH Biomedical Research Centre. Tobias Hauser is supported by a Wellcome Sir Henry Dale Fellowship (211155/Z/18/Z) and a grant from the Jacobs Foundation. Peter Fonagy is in receipt of a National Institute for Health Research (NIHR) Senior Investigator Award (NF-SI-0514-10157). P. Fonagy was in part supported by the NIHR Collaboration for Leadership in Applied Health Research and Care (CLAHRC) North Thames at Barts Health NHS Trust. The views expressed are those of the authors and not necessarily those of the NHS, the NIHR or the Department of Health.

## Competing Interests

E. Bullmore is supported by and holds stock in GlaxoSmithKline Ltd. The other authors have no financial or non-financial competing interests to declare.

