## Supplement (all figures and tables) for "Childhood socio-economic disadvantage predicts reduced myelin growth across adolescence and young adulthood"

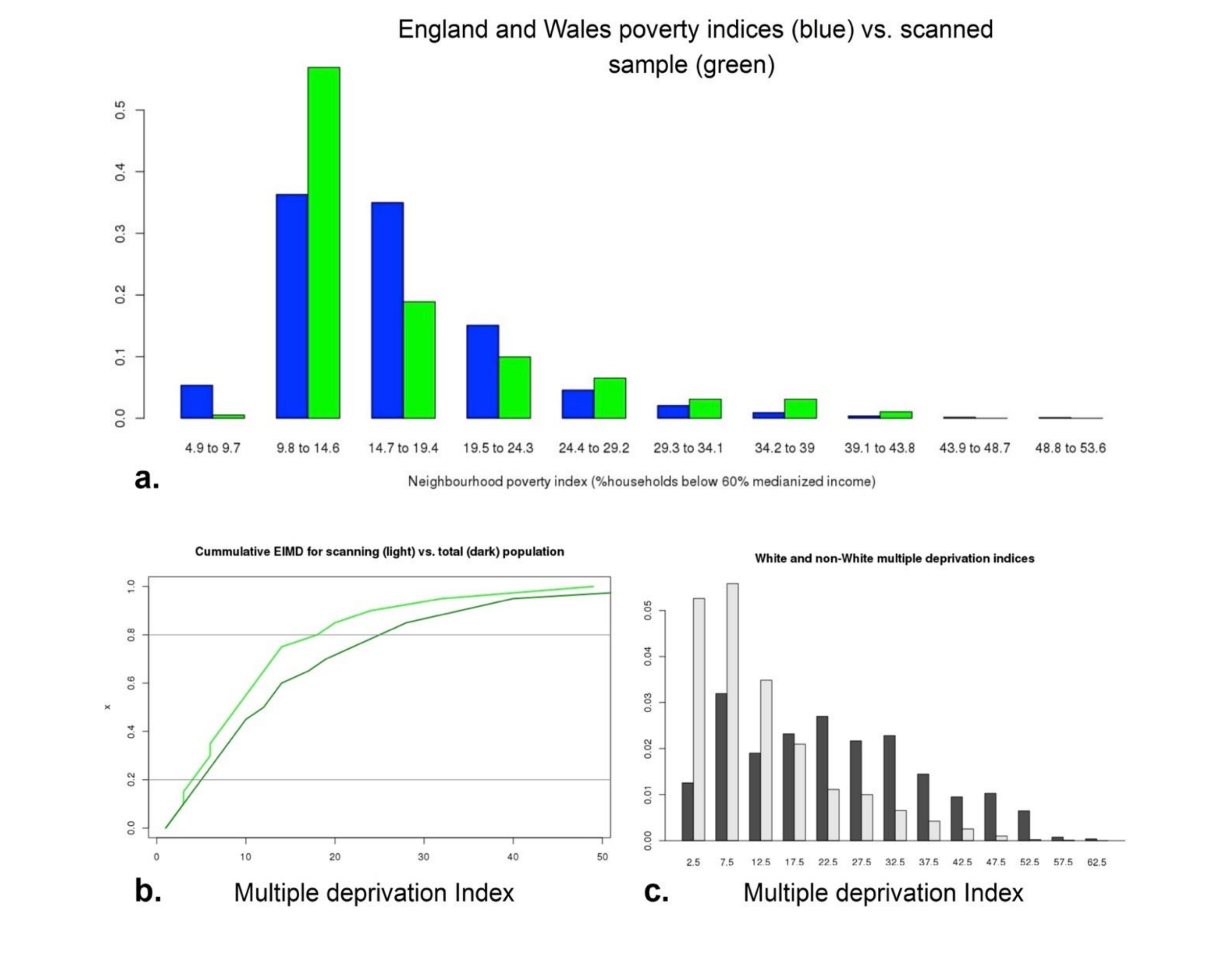


**Figure S1. The scanned sample was approximately representative of the English population with respect to socio-economic deprivation. a.** This sample (in green) contained somewhat fewer worse-deprivation participants than England and Wales as a whole (blue) **b.** Cumulative distribution of SED based on the 2015 English Index of Multiple Deprivation (EIMD), an ONS[10] measure related to the neighbourhood poverty index (Pearson *r*=0.77, p<1e-10). Here we use EIMD as it was available for the entire N=2400 'community sample' from which the scanned sample was recruited. Again, the scanned sample (light green) had a higher density of low disadvantage and lower levels of EIMD, Wilcoxon p=1.7e-7. The mean mood score of the scanned was also better than that of the non-scanned, but the expected relationship between increasing SED and worse mood was similar in the scanned and non-scanned samples (Pearson correlation 0.15 and 0.09, uncorr. *p* 0.01 and 0.0003 respectively). Similarly, the relationship between IQ and SED was similar, and significant, in the scanned sample compared to a non-scanned (but still volunteering to attend the laboratory), N=486 sub-sample of the non-scanned group (Pearson r = -0.13 for both, p=0.0012 uncorr. for these two groups together). **c.** The role of ethnicity. Whites (light grey) were economically advantaged compared to non-whites (dark bars). In the community sample, non-whites also had greatly increased SED compared to whites (EIMD: Wilcoxon test $p < 1e-15$).

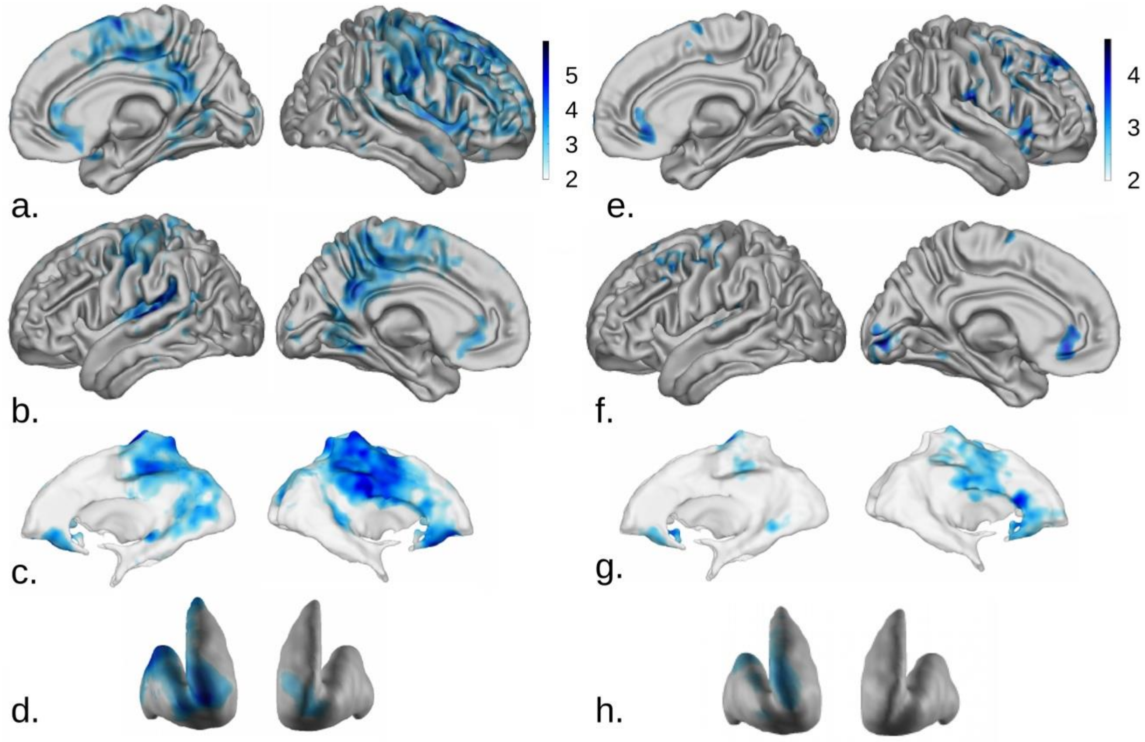


**Figure S2. Exposure to SED during childhood accounts for slowed down myelin marker growth more robustly than currently living in a poor neighbourhood. a.-d**. Similar to Figure 1, areas where MT growth is slower when assessed longitudinally over visits (N=328/185 scans/subjects), as a function of reporting living in a deprived area before age 12. More explanations follow below. **e.-h.** As per a.-d., but the independent variable is deprivation status of the current neighbourhood of residence, on the same sample of participants, i.e. 'current SED'. Analysis of the entire N=479/288 scans/subjects sample strongly resembles e.-h. Rows show grey matter in right hemisphere, then left hemisphere, then core white matter, then striatal grey matter. Only the areas of Figure 1 Where current SED shows some significant associations at the more lenient 15% FDR level are shown. All but 40 in this n=185 participant sample did not change their level of neighbourhood deprivation between before age 12 and testing; therefore we can say with some confidence that exposure during early development is associated with changes in MT growth (a.-d.) but the low economic mobility implies that the (lack of) impact of current deprivation has to be interpreted with greater caution.

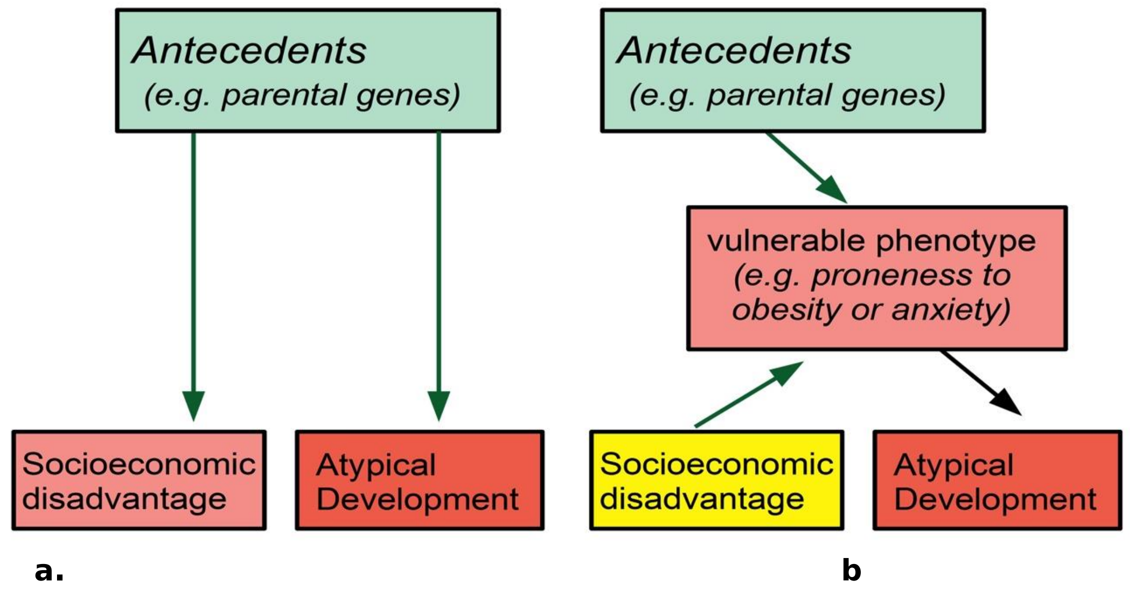


**Figure S35. Psycho-biological approaches to SED-induced effects on brain growth. (a)** Horizontal pleiotropy: An early antecedent causes atypical development side-by-side with another phenotype (here, SED). Hypothetically, genes conferring suboptimal neurocognition in the parents cause them to fail economically, while independently and deterministically give rise to atypical myelination in their offspring. **(b)** Vertical pleiotropy: An antecedent gives rise to a 'vulnerable phenotype', which precedes the second, observed phenotype (in our example, atypical myelination). SED amplifies progression through this vertical cascade, whereas a health-promoting socio-economic status would block it. Hypothetically, genes conferring a ‘sweet tooth’, or high neuroticism may affect neurodevelopment only through obesity, or chronic anxiety. However, the cascade is realized only if SED contributes to poor nutrition, or chronic stress. Horizontal pleiotropy is too restrictive in the light of our results, but vertical pleiotropy is more likely. Vertical and horizontal pleiotropy usually refer to genetic early antecedents, but in our case other early developmental insults might also initiate vulnerable phenotypes.

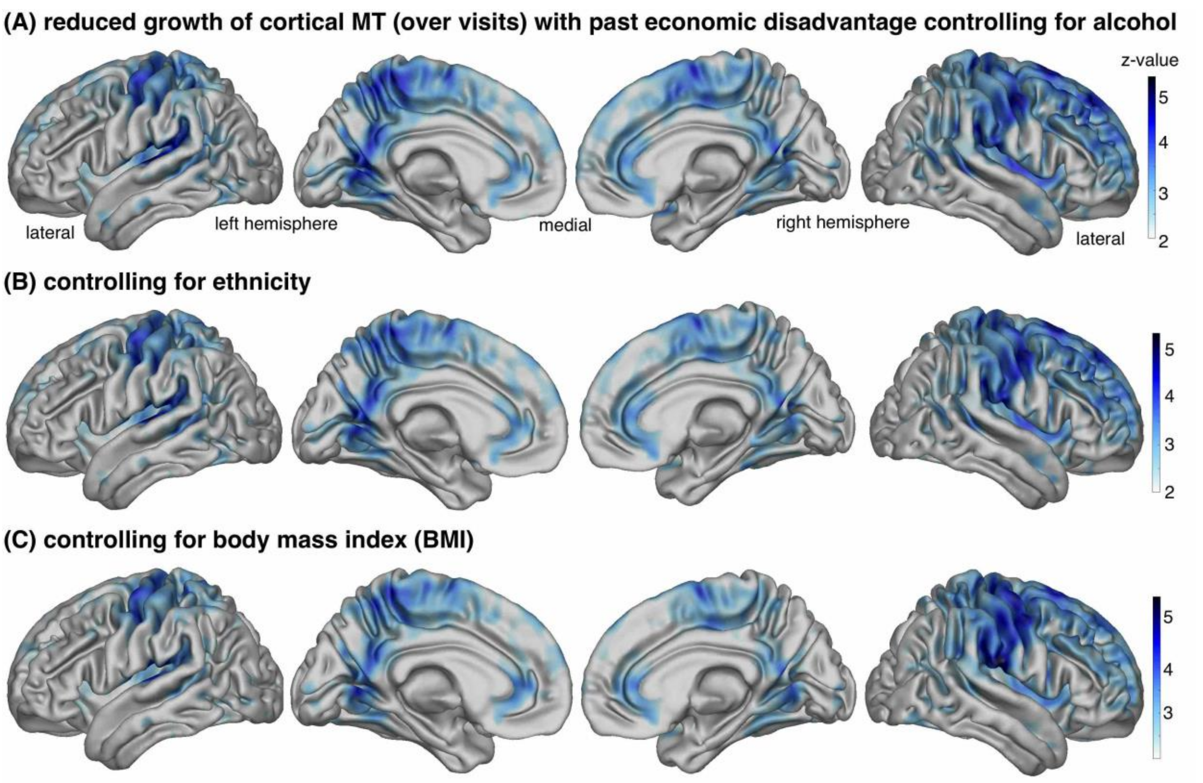


**Figure S4**. **Slower growth of myelin-sensitive MT as early life SED increases is not explained by alcohol, ethnicity or body mass index (BMI).** We present z-maps showing negative SED by time/visit interactions, $p<.05$ FDR corrected, N=328/185 scans/subjects, 45.7% female, when additionally controlling for multiple covariates and their respective time/visit interactions in A-C).

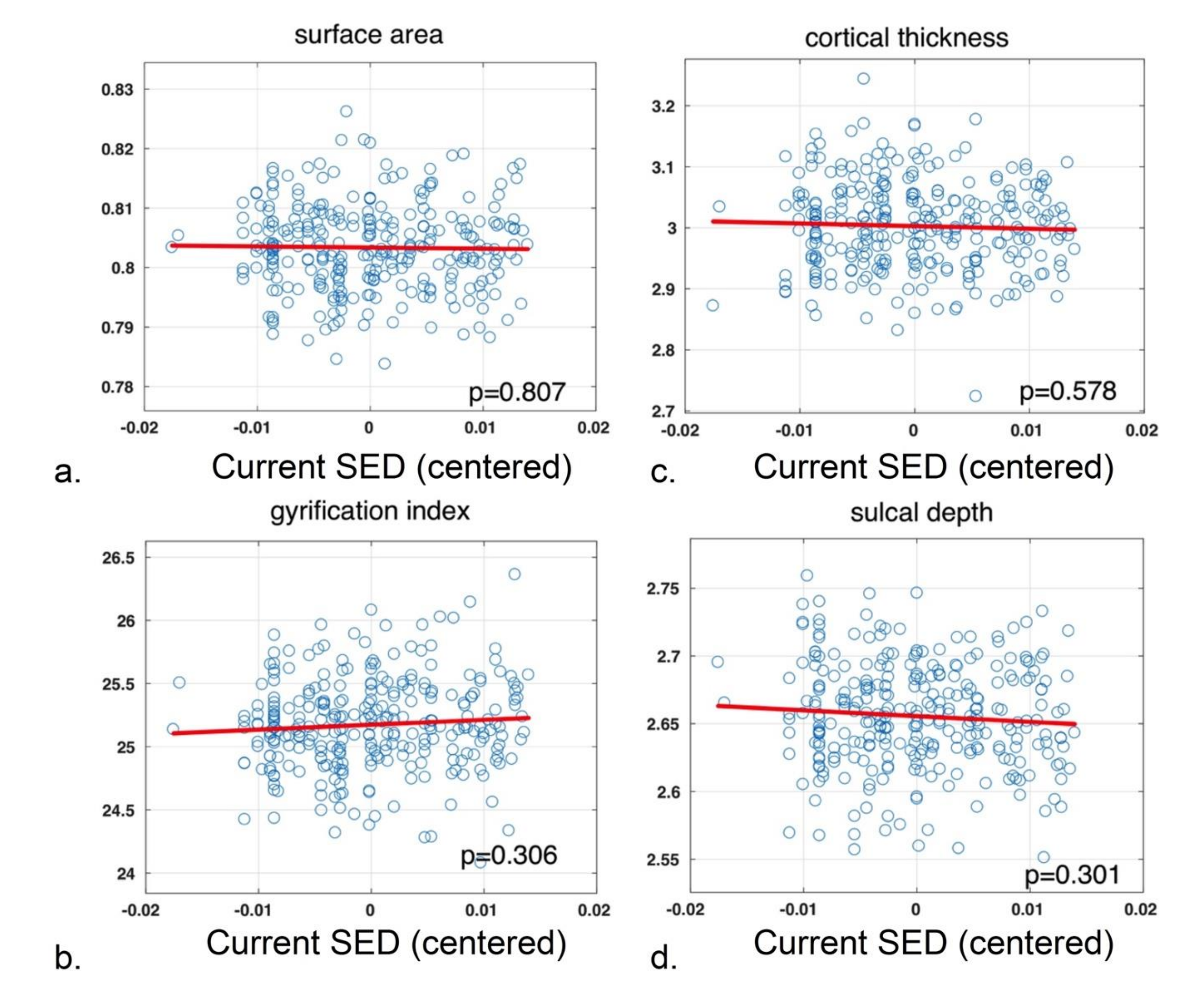


**Figure S5. Morphometric analysis in relation SED.** Measures, including some previously found to correlate with measures of SED different to the ones in our study, were not associated with socio-economic disadvantage. All global surface-based analyses shown here are cross-sectional and controlled for age, sex and their interaction. Linear fits are shown, but quadratic components were not significant either. **a.** to **d.** show representative analyses where global morphometric measures are plotted against current SED. Very similar, non-significant results were found when global morphometric measures were tested against early SED. Additionally, longitudinal analysis of local grey matter volume (GMV) in exact parallel to the main MT analyses showed no evidence for SED-related modulation of GMV shrinkage over visits. This suggests that MT saturation reveals a specific impairment of myelin growth trajectories.

### Supplementary Table 1. SPM results for negative early life SED by time/visit interaction on myelin-sensitive MT within grey matter

| brain region | cluster size | p(FDR) voxelwise | Z | p(unc) | x | y | z |
|---|---|---|---|---|---|---|---|
| **cortical gray matter** | | | | | | | |
| right posterior superior frontal gyrus | 367123 | 9.69E-04 | 5.47 | 2.2E-08 | 13 | 13 | 68 |
| left central operculum | | 9.69E-04 | 5.12 | 1.5E-07 | -52 | -17 | 12 |
| right central operculum/insula | | 9.69E-04 | 5.03 | 2.4E-07 | 39 | 0 | 13 |
| right precentral gyrus | | 9.69E-04 | 4.99 | 3.1E-07 | 51 | -6 | 49 |
| left medial postcentral gyrus | | 9.69E-04 | 4.89 | 5.1E-07 | -9 | -43 | 64 |
| left superior parietal lobe | | 9.69E-04 | 4.68 | 1.4E-06 | -35 | -49 | 52 |
| right postcentral gyrus | | 9.69E-04 | 4.55 | 2.6E-06 | 22 | -32 | 71 |
| left precentral gyrus | | 9.69E-04 | 4.53 | 3.0E-06 | -39 | -16 | 42 |
| right medial orbital gyrus | 1388 | 0.0029 | 3.59 | 1.6E-04 | 13 | 32 | -17 |
| right inferior occipital gyrus | 508 | 0.0064 | 3.16 | 7.9E-04 | 34 | -76 | 6 |
| left medial orbital gyrus | 535 | 0.012 | 2.77 | 0.003 | -19 | 12 | -15 |
| subcortical and cerebellar gray matter | | | | | | | |
| right inf. post. cereb. lobule VIIIB | 66522 | 0.003 | 5.45 | 2.5E-08 | 14 | -41 | -54 |
| left inf. post. cereb. lobule X | | 0.003 | 4.64 | 1.7E-06 | -16 | -38 | -45 |
| left inf. post. cereb. lobule VIIIA | | 0.003 | 4.62 | 1.9E-06 | -18 | -61 | -56 |
| right ant. cereb. lobule III | | 0.003 | 4.43 | 4.7E-06 | 5 | -42 | -12 |
| left sup. post. cereb. lobule VI | | 0.003 | 4.19 | 1.4E-05 | -22 | -54 | -28 |
| left sup. post. cereb. lobule crus II | | 0.003 | 4.16 | 1.6E-05 | -51 | -51 | -43 |
| right ant. cereb. lobule IV | | 0.004 | 3.95 | 3.9E-05 | 24 | -31 | -26 |
| right inf. post. cereb. lobule VIIIA | | 0.004 | 3.84 | 6.1E-05 | 39 | -51 | -56 |
| right ant. caudate | 4987 | 0.004 | 3.78 | 7.9E-05 | 16 | 28 | -4 |
| right putamen | | 0.005 | 3.75 | 8.7E-05 | 29 | -1 | 14 |

SPM longitudinal SwE results table testing for negative early life SED by time/visit interaction effects on MT in cortical and subcortical grey matter accounting for covariates and confounds (cf. methods and supplementary notes on modelling, n=328/185 scans/subjects). Voxel resolution 1mm isotropic. Voxelwise FDR corrected ($p<0.05$) reporting peaks and clusters with up to 8 local maxima more than 24 mm apart, applied extent threshold k=500 voxel. We report FDR on whole-brain level. Using SwE covariance type 'classic', effective degrees of freedom per subject were estimated as 0.973.

**Supplementary Table 2. SPM results for negative early life SED by time/visit interaction on myelin-sensitive MT within white matter**

| Brain region | cluster size | p(FDR) voxelwise | Z | p(unc) | x | y | z |
|---|---|---|---|---|---|---|---|
| **cortex-adjacent white matter** | | | | | | | |
| right postcentral gyrus | 73047 | 0.014 | 4.40 | 5.5E-06 | 47 | -14 | 37 |
| right medial precentral gyrus | | 0.014 | 4.25 | 1.1E-05 | 15 | -10 | 46 |
| right supplementary motor area | | 0.014 | 4.18 | 1.5E-05 | 7 | -2 | 68 |
| right medial orbital gyrus | | 0.014 | 3.95 | 3.9E-05 | 19 | 34 | -7 |
| left precuneus | 39333 | 0.014 | 4.01 | 3.0E-05 | -5 | -58 | 57 |
| left superior postcentral gyrus | | 0.014 | 3.95 | 4.0E-05 | -17 | -41 | 70 |
| left inf. postcentral gyrus | | 0.014 | 3.86 | 5.6E-05 | -47 | -23 | 46 |
| left sup. temporal gyrus | | 0.014 | 3.64 | 1.3E-04 | -56 | -40 | 17 |
| right precuneus | 1407 | 0.014 | 3.86 | 5.6E-05 | 5 | -55 | 17 |
| right ant. middle frontal gyrus | 660 | 0.014 | 3.78 | 7.7E-05 | 36 | 57 | 9 |
| right post. middle frontal gyrus | | 0.014 | 3.31 | 4.7E-04 | 35 | 38 | 35 |
| left post. cingulate gyrus | 2556 | 0.014 | 3.77 | 8.0E-05 | -5 | -49 | 26 |
| left lingual gyrus | 715 | 0.014 | 3.54 | 2.0E-04 | -10 | -60 | 4 |
| right ant. sup. frontal gyrus | 1357 | 0.014 | 3.53 | 2.1E-04 | 24 | 62 | 12 |
| right post. sup. frontal gyrus | | 0.015 | 2.98 | 0.0015 | 7 | 47 | 45 |
| right lingual gyrus | 756 | 0.018 | 2.84 | 0.0023 | 7 | -61 | 6 |
| white matter core areas | | | | | | | |
| rigth ant. corona radiata | 26531 | 0.01 | 3.98 | 3.4E-05 | 18 | 33 | -5 |
| right sup. corona radiata | | 0.01 | 3.94 | 4.1E-05 | 19 | -12 | 44 |
| right external capsule | | 0.01 | 3.87 | 5.5E-05 | 31 | 2 | 10 |
| left post. thalamic radiation | 4465 | 0.011 | 3.16 | 7.8E-04 | -41 | -42 | 2 |
| left sup. longitudinal fascic. | 1527 | 0.013 | 2.98 | 0.0014 | -38 | -24 | 28 |

SPM longitudinal SwE results table testing for negative early life SED by time/visit interaction effects on MT in white matter accounting for covariates and confounds (cf. methods and supplementary notes on modelling, n=328/185 scans/subjects). Voxel resolution 1mm isotropic. Voxelwise FDR corrected ($p<0.05$) reporting peaks and clusters with up to 4 local maxima more than 24 mm apart, applied extent threshold k=500 voxel. We report FDR on whole-brain level. Using SwE covariance type ‘classic’, effective degrees of freedom per subject were estimated as 0.973.

**Supplementary Table 3. SPM results for early life SED by parenting quality by time/visit interaction on cortical myelin-sensitive MT**

| Brain region | cluster size | p(FDR) voxelwise | Z | p(unc) | x | y | z |
|---|---|---|---|---|---|---|---|
| **cortical gray matter** | | | | | | | |
| right ant. middle frontal gyrus | 5327 | 2.9E-04 | 5.69 | 6.5E-09 | 29 | 55 | -3 |
| right ant. sup. frontal gyrus | | 0.011 | 4.08 | 2.2E-05 | 27 | 52 | 36 |
| left inf. frontal angular gyrus | 10028 | 0.006 | 4.33 | 7.4E-06 | -49 | 33 | -8 |
| left ant. middle frontal gyrus | | 0.007 | 4.27 | 9.6E-06 | -43 | 49 | 19 |
| left inf. frontal gyrus | | 0.007 | 4.25 | 1.0E-05 | -54 | 25 | 20 |
| left ant. middle frontal gyrus | | 0.018 | 3.77 | 8.3E-05 | -28 | 56 | -3 |
| right ant. middle frontal gyrus | 426 | 0.007 | 4.24 | 1.1E-05 | 34 | 52 | 11 |
| left medial superior frontal gyrus | 335 | 0.013 | 4.00 | 3.1E-05 | -6 | 55 | 30 |
| right post. middle frontal gyrus | 413 | 0.021 | 3.70 | 1.1E-04 | 36 | 8 | 55 |
| left sup. frontal gyrus | 301 | 0.023 | 3.61 | 1.5E-04 | -16 | 28 | 55 |
| **cortex-adjacent white matter** | | | | | | | |
| right ant. middle frontal gyrus | 588 | 0.017 | 4.65 | 1.6E-06 | 32 | 56 | -2 |
| right sup. frontal gyrus | 295 | 0.017 | 4.44 | 4.4E-06 | 14 | 60 | -8 |

SPM longitudinal SwE results table testing for positive early life SED by parenting quality by time/visit interaction effects on MT in cortical grey and adjacent white matter accounting for covariates and confounds (cf. methods and supplementary notes on modelling, n=328/185 scans/subjects). Voxel resolution 1mm isotropic. Voxelwise FDR corrected ($p<0.05$) reporting peaks and clusters with up to 4 local maxima more than 24 mm apart, applied extent threshold k=250 voxel. Using SwE covariance type ‘classic’, effective degrees of freedom per subject were estimated as 0.9568.

**Supplementary Table 4. SPM results for parenting quality by sex interaction on cortical myelin-sensitive MT**

| Brain region | cluster size | p(FDR) voxelwise | Z | p(unc) | x | y | z |
|---|---|---|---|---|---|---|---|
| **cortical gray matter** | | | | | | | |
| right precuneus | 94498 | 0.003 | 5.30 | 5.7E-08 | 7 | -52 | 23 |
| right superior parietal lobe | | 0.003 | 5.14 | 1.3E-07 | 31 | -53 | 48 |
| right post. cingulate gyrus | | 0.003 | 5.12 | 1.5E-07 | 3 | -38 | 45 |
| left precentral gyrus | 7290 | 0.003 | 5.10 | 1.7E-07 | -41 | -12 | 32 |
| left postcentral gyrus | | 0.004 | 4.05 | 2.6E-05 | -24 | -28 | 56 |
| right precentral gyrus | 15031 | 0.003 | 5.03 | 2.4E-07 | 34 | -15 | 43 |
| right postcentral gyrus | | 0.003 | 4.21 | 1.3E-05 | 27 | -36 | 53 |
| right supramarginal gyrus | | 0.004 | 4.07 | 2.4E-05 | 60 | -18 | 42 |
| left inf. temporal gyrus | 4774 | 0.003 | 4.62 | 1.9E-06 | -53 | -47 | -26 |
| left middle temporal gyrus | | 0.006 | 3.77 | 8.2E-05 | -47 | 3 | -27 |
| left middle temporal gyrus | | 0.008 | 3.56 | 1.9E-04 | -60 | -25 | -10 |
| left post. superior frontal gyrus | 3450 | 0.003 | 4.55 | 2.7E-06 | -21 | -10 | 58 |
| left middle frontal gyrus | | 0.009 | 3.40 | 3.3E-04 | -26 | 14 | 53 |
| right central operculum | 4475 | 0.003 | 4.30 | 8.7E-06 | 42 | -10 | 17 |
| right parietal operculum | | 0.005 | 3.82 | 6.6E-05 | 57 | -28 | 23 |
| left medial orbital gyrus | 3470 | 0.003 | 4.22 | 1.2E-05 | -24 | 31 | -15 |
| right post. gyrus rectus | 6722 | 0.004 | 4.10 | 2.1E-05 | 6 | 25 | -22 |
| right ant. gyrus rectus | 3600 | 0.007 | 3.67 | 1.2E-04 | 3 | 62 | -22 |

SPM longitudinal SwE results table testing for positive parenting quality by sex interaction effects on MT in cortical grey matter accounting for covariates and confounds (cf. methods and supplementary notes on modelling, n=328/185 scans/subjects). Voxel resolution 1mm isotropic. Voxelwise FDR corrected ($p<0.05$) reporting peaks and clusters with up to 3 local maxima more than 24 mm apart, applied extent threshold k=3000 voxel. We report FDR on whole-brain level. Using SwE covariance type ‘classic’, effective degrees of freedom per subject were estimated as 0.9568.

**Supplementary Table 5. SPM results for body mass index effects on cortical myelin-sensitive MT**

| Brain region | cluster size | p(FDR) voxelwise | Z | p(unc) | x | y | z |
|---|---|---|---|---|---|---|---|
| **cortical gray matter** | | | | | | | |
| right frontal operculum | 447 | 0.021 | 5.06 | 2.1E-07 | 29 | 23 | 13 |
| left superior temporal gyrus | 385 | 0.021 | 4.72 | 1.2E-06 | -48 | -4 | -16 |
| right subgenual gyrus | 1256 | 0.021 | 4.54 | 2.8E-06 | 9 | 37 | -3 |
| right ant. midcingulate gyrus | | 0.029 | 4.04 | 2.7E-05 | 2 | 8 | 44 |
| right ant. cingulate gyrus | | 0.034 | 3.89 | 5.0E-05 | 4 | 27 | 24 |
| left sup. frontal gyrus | 250 | 0.027 | 4.08 | 2.2E-05 | -24 | -9 | 54 |

SPM longitudinal SwE results table testing for negative body mass index effects on MT in cortical grey matter accounting for covariates and confounds (cf. methods and supplementary notes on modelling, n=277/155 scans/subjects). Voxel resolution 1mm isotropic. Voxelwise FDR corrected ($p<0.05$) reporting peaks and clusters with up to 3 local maxima more than 24 mm apart, applied extent threshold k=250 voxel. We report FDR on whole-brain level. Using SwE covariance type ‘classic’, effective degrees of freedom per subject were estimated as 0.9484.

**Supplementary table 6.** Neuroscience in Psychiatry consortium author list.

| Neuroscience in Psychiatry Network Study & Consortium Author list | |
|---|---|
| Principal Investigators | Edward Bullmore (CI from 01/01/17) |
| | Ian Goodyer (CI until 01/01/2017) |
| | Raymond Dolan |
| | Peter Fonagy |
| | Peter Jones |
| NSPN funded staff | Michael Moutoussis |
| | Tobias U. Hauser |
| | Petra Vértes |
| | Kirstie Whitaker |
| | Gita Prabhu |
| | Laura Villis |
| | Junaid Bhatti |
| | Becky Inkster |
| | Cinly Ooi |
| | Barry Widmer |
| | Ayesha Alrumaithi |
| | Sarah Birt |
| | Kalia Cleridou |
| | Hina Dadabhoy |
| | Sian Granville |
| | Elizabeth Harding |
| | Alexandra Hopkins |
| | Daniel Isaacs |
| | Janchai King |
| | Danae Kokorikou |
| | Harriet Mills |
| | Ciara O'Donnell |
| | Sara Pantaleone |
| Affiliated Scientists | Pasco Fearon |
| | Anne-Laura van Harmelen |
| | Rogier Kievit |
